# Glial cell and perineuronal net interactions in the dorsal striatum of aged mice

**DOI:** 10.64898/2026.03.25.714174

**Authors:** Zachary A Colon, Alejandro Gamboa Fuentes, Scott Litwiler, Kathleen A Maguire-Zeiss

## Abstract

Elucidating how normal aging increases vulnerability to neurodegeneration remains a major gap in our understanding of disease risk and progression. The dorsal striatum serves as the primary input nucleus of the basal ganglia and is a key region implicated in multiple neurodegenerative diseases (NDDs) (1). In Colon et al. 2025 (2), we examined the impact of normal aging on neuroinflammatory signaling and perineuronal net (PNN) homeostasis within the dorsal striatum. We observed age-associated shifts in the inflammatory landscape and evidence of increased microglial activation, yet PNN homeostasis was largely preserved (2). PNNs are highly organized extracellular matrix (ECM) specializations that preferentially enwrap the soma and proximal dendrites of fast-spiking GABAergic parvalbumin (PV) interneurons, where they contribute to the regulation of synaptic plasticity and provide protection against oxidative stress (3,4). Building on these findings, we developed a working hypothesis to explain the apparent preservation of PNN homeostasis despite an aging-associated pro-inflammatory environment.

The shift toward a pro-inflammatory milieu, together with increased gliosis and phagocytic activity, would be expected to impact the maintenance and integrity of perineuronal nets. The observed increase in phagocytosis-related markers may reflect microglia-directed activity as well as contributions from additional central nervous system (CNS) cell populations. Microglia are specialized embryonic-derived myeloid cells that serve as the resident immune cells of the brain and contribute to PNN homeostasis under physiological conditions (5). In Colon et al. 2025, we observed evidence of microgliosis (e.g., morphological changes, *Iba1, Trem2*) along with elevated expression of markers associated with phagocytosis (e.g., *Cd68*) and extracellular matrix–modifying proteases (e.g., *Mmp9, Adam17*) capable of cleaving key PNN components (2). Importantly, *Cd68* expression is not exclusive to microglia and has been detected in brain infiltrating macrophages, reactive astrocytes, and neutrophils during inflammation (6–8). Thus, increased *Cd68* levels may not solely reflect microglial phagocytic activation but may also reflect astrocyte reactivity and phagocytic phenotypes. Furthermore, astrocytes are the most abundant glial cell in the brain, and they play a major role in maintaining CNS homeostasis by regulating extracellular neurotransmitter concentrations, providing metabolic support, contributing to the synthesis and remodeling of PNN components, and modulating neuronal communication through their involvement in the tetrapartite synapse (9–12). Astrocytes can also phagocytosis microglial debris, myelin, and synapses (7).

To better define the cellular source of phagocytic activity and its relationship to PNN remodeling in aging, we performed immunostaining for microglia (Iba1^+^), astrocytes (GFAP^+^), phagolysosomal activity (CD68^+^), and PNNs using *Wisteria floribunda* agglutinin (WFA^+^), enabling us to assess the spatial relationship between phagocytosis and PNN components.

## Materials & Methods

Wild-type C57BL/6 mice (n = 5/group) were aged to 4 months (Young) or 22 months (Aged) and subjected to a battery of behavioral assessments prior to tissue collection for downstream RNA and immunohistochemical (IHC) analyses (previously published, Colon et al. 2025 (2)). Figure 1 outlines experimental details pertinent to this commentary. Coronal 30-µm hemisections containing the dorsal striatum were processed in two independent staining/imaging cohorts: (1) microglia were labeled with Iba1 (rabbit; FUJIFILM Wako, #P3088) in combination with the lysosomal/phagocytic marker CD68 (rat; Bio-Rad, #MCA1957) and perineuronal nets (PNNs) were visualized using fluorescein-conjugated *Wisteria floribunda* agglutinin (WFA; Vector Laboratories, #FL-1351); (2) astrocytes were labeled with GFAP (rabbit; Cell Signaling Technology, #12389) together with WFA to visualize PNNs. Secondary antibodies included goat anti-rat Alexa Fluor 594 (Invitrogen, #A48264), goat anti-rabbit Alexa Fluor 635 (Invitrogen, #A31577), and goat anti-mouse Alexa Fluor 594 (Invitrogen, #A21125), as appropriate for each experiment. Quantification of total cell number, fluorescence intensity, and regression analyses were performed across the entire dorsal striatum. Single-cell analyses were conducted using high-magnification z-stack images (Zeiss Axio Imager.Z2 microscope) acquired from randomly selected WFA+ PNNs, with all fully captured glial cells within each field included for analysis. All imaging and quantification were performed by investigators blinded to experimental group (see Colon et al. 2025 (2) for details).

**Figure 1:**
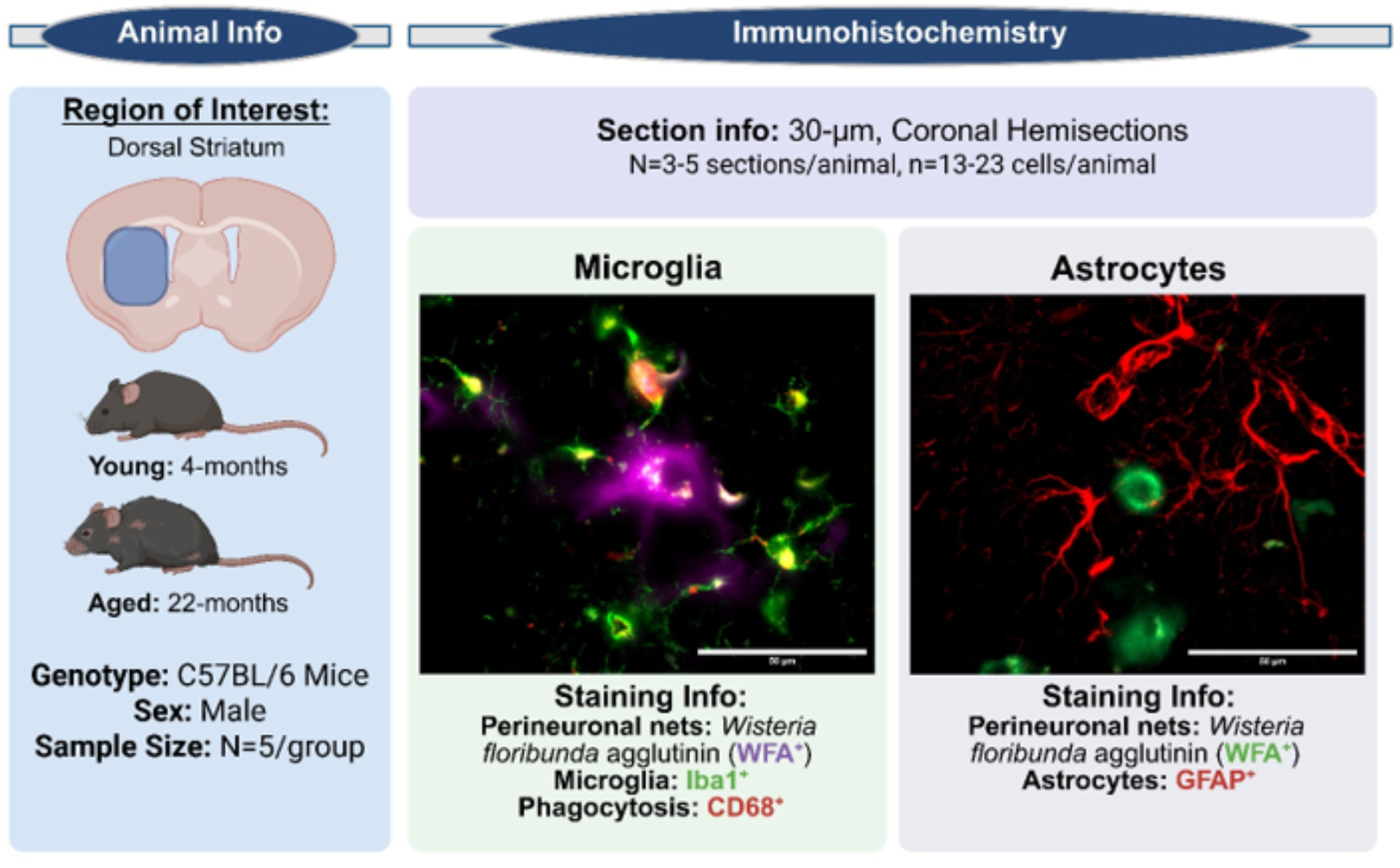
Experimental Design.

## Results

### Normal aging increases the expression of CD68^+^ in microglia

Microglia within the aged dorsal striatum displayed canonical morphological features consistent with microgliosis, accompanied by elevated expression of *Cd68* (a lysosomal/phagocytic marker) and *Mmp9* (matrix metalloproteinase-9; a protease capable of degrading extracellular matrix and PNN components) (2). Given that increased phagocytic capacity and proteolytic activity would be expected to compromise PNN integrity, the observation that overall PNN homeostasis was preserved suggests that these age-associated changes may not directly translate to enhanced PNN degradation, or alternatively that compensatory mechanisms maintain net stability despite heightened microglial activation (2).

To investigate microglia at the whole-region level, we detected no significant age-related differences in total microglial abundance (Fig. 2A) within the dorsal striatum. However, single-cell analyses revealed significantly increased Iba1^+^ and CD68+ immunoreactivity per microglia in the Aged animals (Fig. 2B–C), indicating an age-dependent shift toward a more activated and lysosome-enriched microglial phenotype without gross changes in cell number. Additionally, Aged mice exhibited an upward trend in WFA^+^ signal per microglia (Fig. 2D). Regression analyses demonstrated a significant positive relationship between Iba1^+^ and CD68^+^ expression in both Young and Aged animals (Fig. 2E), supporting the interpretation that increased CD68 levels are coupled to microglial activation state. In contrast, no age-dependent association was observed between Iba1^+^ and WFA^+^ nor between CD68^+^ and WFA^+^ (Fig. 2F–G), suggesting that WFA-labeled PNN-associated glycans are not robustly linked to microglial phagolysosomal content under aging conditions. This raises the possibility that microglial engulfment may preferentially involve non-glycan PNN constituents, warranting additional analyses using alternative PNN markers.

**Figure 2:**
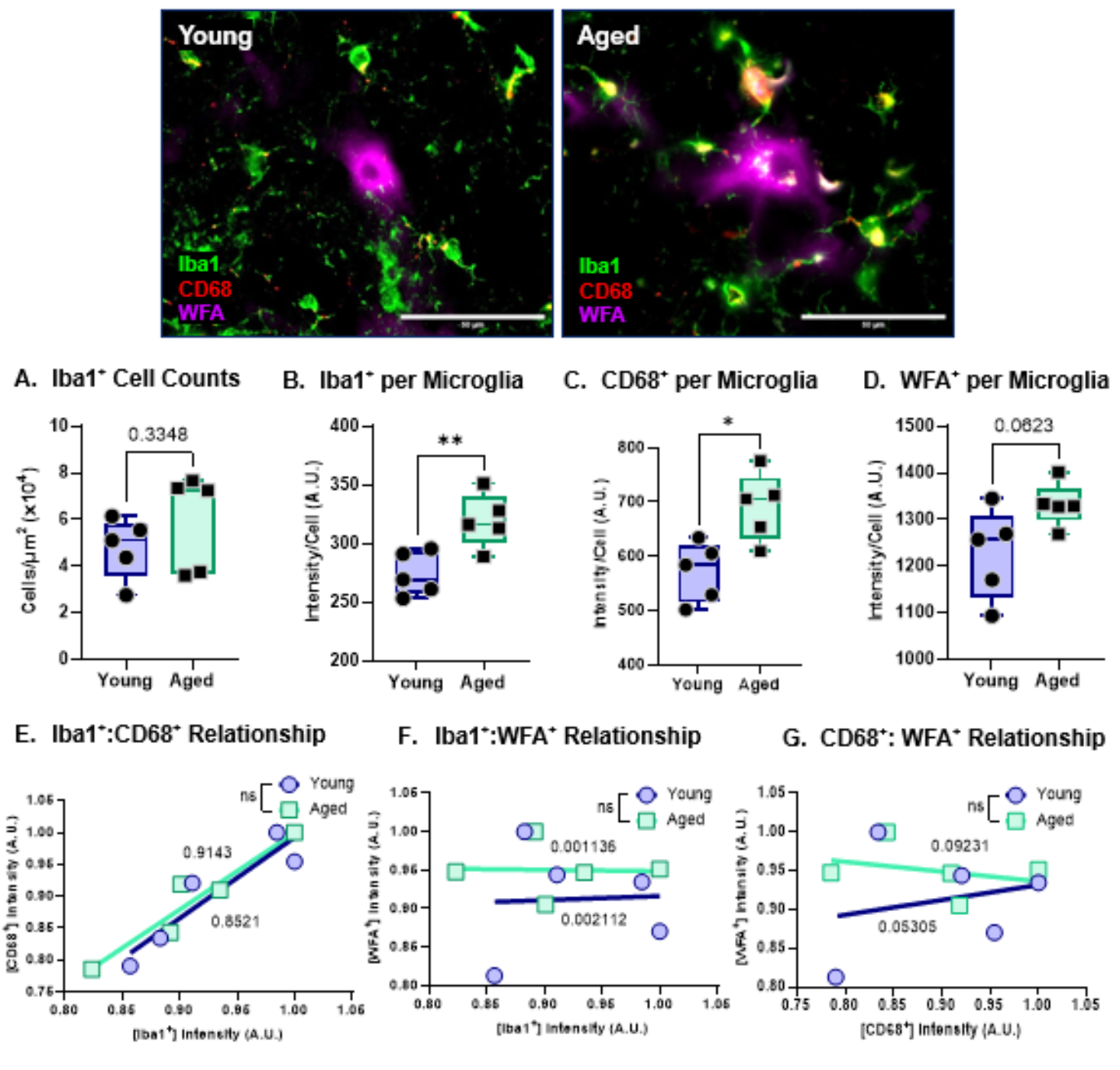
Normal aging promotes microglial activation and increased CD68^+^expression in the dorsal striatum. A Quantification of Iba1^+^ cell counts for the total striatum. Data was normalized to area. B. Quantification of Iba1^+^ intensity per Iba1^+^ cell. C. Quantification of CD68^+^ intensity per Iba1^+^ cell D. Quantification of WFA^+^ intensity per Iba1^+^ cell. E. Linear regression analysis of Ibal^+^ intensity versus CD68^+^ intensity per cell. F. Linear regression analysis of Ibal^+^ intensity versus UTA^+^ intensity per cell. G. Linear regression analysis of CD68^+^ intensity versus UTA^+^ intensity per cell. A Data represented as mean ± SENL data was tested for normality prior to statistical analysis. Two-tailed t-tests or Welch’s t-test used when appropriate. Regression analysis: R^2^ values of corresponding linear regressions label on graph and t-test comparison of slopes used, p-values: * <0.05, * * <0.01. N - 4-5 mice/group, 3-5 sections/mouse, 13-41 cells/mouse.

Collectively, these findings indicate that normal aging is associated with increased microglial activation and lysosomal/phagocytic marker expression at the single-cell level yet occurs without detectable disruption of overall PNN homeostasis. This dissociation supports the existence of compensatory regulatory processes—potentially including astrocyte-mediated maintenance of extracellular matrix structure—that preserve PNN integrity despite heightened microglial reactivity.

### Normal aging increases astrocyte proliferation and association with perineuronal nets, but astrocytes do not display reactive morphology

To assess age-related changes in astrocytes, we next quantified astrocyte abundance, morphological features, and spatial interactions with perineuronal nets in the dorsal striatum. Transcriptomic analyses revealed increased expression of *Gfap* and *C3*, two markers commonly associated with A1-like reactive astrocytes, raising the possibility that astrocytic reactivity increases during normal aging (2). Consistent with this, aged mice exhibited a significant increase in total astrocyte number (Fig. 3A). However, GFAP^+^ immunoreactivity per astrocyte was not significantly altered (Fig. 3B), suggesting that the elevated *Gfap* signal observed at the RNA level is more likely driven by increased astrocyte abundance rather than enhanced GFAP^+^ expression on a per-cell basis.

**Figure 3.**
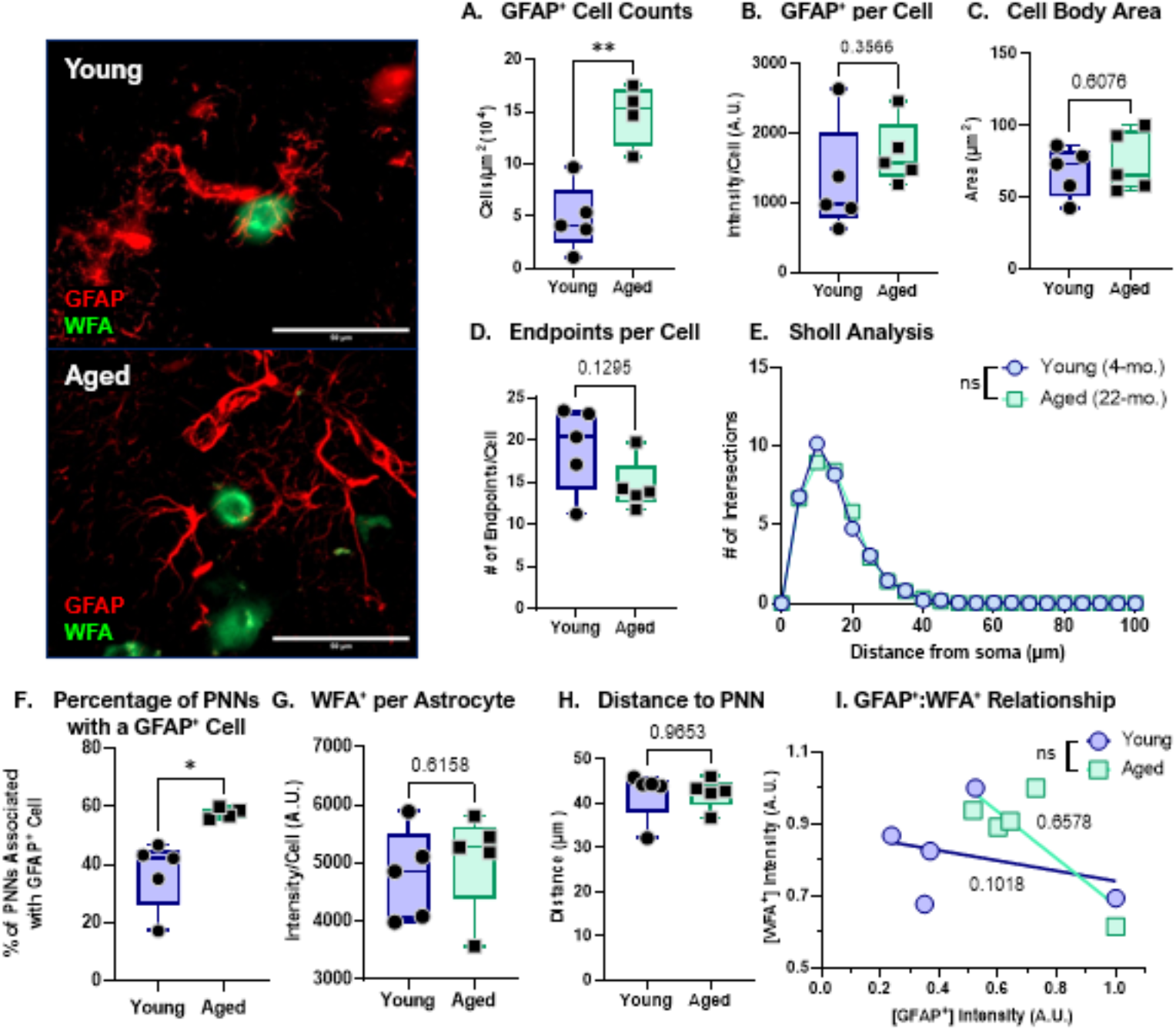
Legend: Normal aging increases astrocyte proliferation but does not increase reactive astrocyles. B. Quantification of GFAP^+^ cell counts for the total striatum. Data was normalized to area. C. Quantification of GFAP^+^ intensity per cell. D. Quantification of GFAP^+^ cell bodv area in the striatum. E. Quantification of average endpoints per GFAP^+^ cell in the striatum. F. Sholl analysis of the branch profile (ramification) of GFAP^+^ cells in the striatum. G. Percentage ofPNNs associated with a GFAP^+^ cells. H. Quantification of WTA intensity per GFAP^+^cell. I. Quantification of distance of GFAP^+^cells to soma of PNN-ensheathed neurons. J. Linear regression analysis of GFAP intensity versus WFA^+^ intensity per cell. R^2^ values of corresponding linear regressions label on graph. Data represented as mean ± SEM, data was tested for normality prior to statistical analysis. Two-tailed t-tests or Welch’s t-test used when appropriate. Sholl analysis: Two-Way ANOVA with Tukey. Regression analysis: t-test comparison of slopes, p-values: * <0.05, ** <0.01. N = 4-5 mice/group, 3-5 sections/mouse, 13-41 cells/mouse.

To further evaluate astrocyte reactivity at the cellular level, we performed morphological analyses of astrocytes in proximity to PNNs. Reactive astrocytes are typically characterized by hypertrophy, including increased soma size and altered process architecture (e.g., shorter, thicker ramifications) (13). In our dataset, soma cell body area did not differ between the Young and Aged groups (Fig. 3C). Similarly, measures of process complexity showed no age-related changes, including the number of endpoints per cell (Fig. 3D) and process length/distribution (Fig. 3E). Collectively, these findings support the interpretation that astrocytes increase in number during aging without exhibiting overt morphological hallmarks of reactivity, consistent with preservation of homeostatic astrocyte function in the aged striatum.

To define how astrocytes interact with PNNs during aging, we next quantified astrocyte–PNN spatial relationships. Regions of interest were generated around individual WFA^+^ PNNs throughout the dorsal striatum, and the proportion of PNNs closely associated with a GFAP^+^ astrocyte was calculated. Aged mice displayed a significantly higher percentage of PNNs associated with astrocytes (Fig. 3F), indicating increased astrocytic coverage or recruitment to PNN-bearing neurons with age. This increased association was not accompanied by changes in WFA^+^ intensity within individual astrocytes (Fig. 3G), astrocyte distance to the PNN-bearing neuronal soma (Fig. 3H), or the degree of association of GFAP^+^ to WFA^+^ signal (Fig. 3I). These data suggest that astrocytes are in close proximity to PNNs and that aging increases the number of astrocytes available to interact with these structures. Together, these findings support a model in which normal aging increases astrocyte abundance. This astrocytic expansion may serve as a compensatory mechanism that buffers the extracellular matrix against age-related increases in microglial activation, phagocytic signaling, and proteolytic pressure, thereby contributing to the maintenance of PNN homeostasis.

## Discussion

Perineuronal net homeostasis is a dynamic process that requires coordination between neurons, microglia, and astrocytes (11). Microglia and astrocytes coordinate PNN regulation through bidirectional signaling that controls ECM production, remodeling, and degradation around PV interneurons (11). Under homeostatic conditions, astrocytes support PNN maintenance by providing metabolic support and secreting key ECM components such as hyaluronan-associated chondroitin sulfate proteoglycans (CSPGs) and link proteins that stabilize net architecture (11,14). In parallel, microglia remain in a surveillant phenotype and help preserve circuit integrity through trophic support and limited remodeling. During aging, microglial activation increases the release of cytokines and chemokines (e.g., IL-6, TNF-α) (15), which would be predicted to promote astrocyte reactivity and shifts astrocytic output toward altered CSPG deposition and increased expression of ECM-modifying proteases. Here we report that despite increases inflammation as well as microglial activation and phagocytosis there is no change in the number of PNNs. While CD68^+^ is increased in the aged microglia and we show a trend towards increased PNN/microglia, the possibility remains that microglia are also targeting synapses, myelin debris, and protein aggregates for phagocytosis. Alternatively, the increased CD68/microglia could reflect an intracellular lysosomal buildup within microglia.

The exact function of aged reactive microglia and the coordinated glial communication that acts to balance PNN stabilization and degradation/remodeling remains unclear and warrants further investigation. It is possible that the proliferation of astrocytes in aged mice is a compensatory mechanism working to balance increased microgliosis. Prior work has demonstrated that experimental removal of PNNs can promote compensatory expansion of astrocytic pericellular coverage around PV interneurons without significantly altering synaptic transmission, suggesting that astrocytes may dynamically adapt to changes in the perineuronal extracellular matrix (16). In addition, PV interneurons have exceptionally high metabolic demands, and astrocytes are essential for sustaining their activity through metabolic coupling mechanisms such as the astrocyte–neuron lactate shuttle (17,18). A caveat is that while the use of GFAP is well established measure for astrogliosis, it is a cytoskeletal protein leaving detailed measures on the astrocytic coverage on the soma of PNN-containing cells remains limited.

## Conclusion

Overall, these findings support an age-related shift toward microglia activation without producing overt disruption of PNN homeostasis. At the single-cell level, microglia in Aged mice exhibited increased Iba1^+^ and CD68^+^ expression, consistent with a more activated, lysosome-enriched phenotype. In parallel, aging increased astrocyte abundance and astrocyte–PNN association without clear morphological hallmarks of reactivity, raising the possibility that astrocytes provide a compensatory stabilizing influence that buffers PNN structure against increased microglial inflammatory and proteolytic pressure. Together, these data suggest that PNNs may remain stable under baseline conditions but potentially vulnerable to degradation following a secondary inflammatory insult, highlighting the importance of coordinated glial communication in maintaining extracellular matrix integrity. We are actively continuing to investigate how microglia and astrocytes interact with PNNs across aging and inflammatory conditions, so stay tuned.

## Acknowledgements

We would like to thank the Georgetown University Department of Neuroscience microscopy CORE. We would like to express our gratitude to the Georgetown University Division of Comparative Medicine veterinarians and veterinary technicians for the care of the mice.

## Author Contributions

ZAC and KMZ designing and conceptualization of experiments. SL performed microglia and CD68 experiments and analysis. AGF performed astrocyte and PNN experiments and analysis for total counts and intensity. ZAC performed astrocytic morphological experiments and analysis. ZAC wrote the majority of the commentary with revisions done by ZAC and KMZ. ZAC created figures and images. All authors read, reviewed, and approved the commentary.

## References

1. Steiner H, Tseng K. Handbook of Basal Ganglia Structure and Function. 2010.

2. Colon ZA, Chan SC, Maguire-Zeiss KA. Age-related inflammatory changes and perineuronal net dynamics: implications for aging. J Neuroinflammation. 2025 Nov 18;22(1):274.

3. Bosiacki M, Gąssowska-Dobrowolska M, Kojder K, Fabiańska M, Jeżewski D, Gutowska I, et al. Perineuronal Nets and Their Role in Synaptic Homeostasis. Int J Mol Sci. 2019 Aug 22;20(17):4108.

4. Cabungcal JH, Steullet P, Morishita H, Kraftsik R, Cuenod M, Hensch TK, et al. Perineuronal nets protect fast-spiking interneurons against oxidative stress. Proceedings of the National Academy of Sciences. 2013 May 28;110(22):9130–5.

5. Crapser JD, Arreola MA, Tsourmas KI, Green KN. Microglia as hackers of the matrix: sculpting synapses and the extracellular space. Cell Mol Immunol. 2021 Nov;18(11):2472–88.

6. Chistiakov DA, Killingsworth MC, Myasoedova VA, Orekhov AN, Bobryshev YV. CD68/macrosialin: not just a histochemical marker. Lab Invest. 2017 Jan;97(1):4–13.

7. Morizawa YM, Hirayama Y, Ohno N, Shibata S, Shigetomi E, Sui Y, et al. Reactive astrocytes function as phagocytes after brain ischemia via ABCA1-mediated pathway. Nat Commun. 2017 Jun 22;8:28.

8. Matsumoto H, Kumon Y, Watanabe H, Ohnishi T, Shudou M, Ii C, et al. Antibodies to CD11b, CD68, and lectin label neutrophils rather than microglia in traumatic and ischemic brain lesions. Journal of Neuroscience Research. 2007;85(5):994– 1009.

9. Schousboe A, Bak LK, Waagepetersen HS. Astrocytic Control of Biosynthesis and Turnover of the Neurotransmitters Glutamate and GABA. Front Endocrinol (Lausanne). 2013 Aug 15;4:102.

10. Verkhratsky A, Parpura V, Li B, Scuderi C. Astrocytes: The Housekeepers and Guardians of the CNS. Adv Neurobiol. 2021;26:21–53.

11. Tewari BP, Chaunsali L, Prim CE, Sontheimer H. A glial perspective on the extracellular matrix and perineuronal net remodeling in the central nervous system. Front Cell Neurosci. 2022 Oct 20;16:1022754.

12. Chelini G, Pantazopoulos H, Durning P, Berretta S. The tetrapartite synapse: a key concept in the pathophysiology of schizophrenia. Eur Psychiatry. 2018 Apr;50:60–9.

13. Sun D, Jakobs TC. Structural Remodeling of Astrocytes in the Injured CNS. Neuroscientist. 2012 Dec;18(6):567–88.

14. Oohashi T, Edamatsu M, Bekku Y, Carulli D. The hyaluronan and proteoglycan link proteins: Organizers of the brain extracellular matrix and key molecules for neuronal function and plasticity. Experimental Neurology. 2015 Dec 1;274:134–44.

15. Franceschi C, Garagnani P, Parini P, Giuliani C, Santoro A. Inflammaging: a new immune–metabolic viewpoint for age-related diseases. Nat Rev Endocrinol. 2018 Oct;14(10):576–90.

16. Tewari BP, Woo AM, Prim CE, Chaunsali L, Patel DC, Kimbrough IF, et al. Astrocytes require perineuronal nets to maintain synaptic homeostasis in mice. Nat Neurosci. 2024 Aug;27(8):1475–88.

17. Inan M, Zhao M, Manuszak M, Karakaya C, Rajadhyaksha AM, Pickel VM, et al. Energy deficit in parvalbumin neurons leads to circuit dysfunction, impaired sensory gating and social disability. Neurobiology of Disease. 2016 Sep 1;93:35– 46.

18. Kim Y, Dube SE, Park CB. Brain energy homeostasis: the evolution of the astrocyte-neuron lactate shuttle hypothesis. Korean J Physiol Pharmacol. 2025 Jan 1;29(1):1–8.

